# Using fingerprinting as a testbed for strategies to improve reproducibility of functional connectivity

**DOI:** 10.1101/2024.06.21.599225

**Authors:** Jivesh Ramduny, Tamara Vanderwal, Clare Kelly

## Abstract

Reproducible functional connectivity-based biomarkers have remained elusive despite the promise of deeply phenotyped consortia. An important component of reproducibility is reliability over repeated measures, often measured by the intraclass correlation coefficient (ICC). Here, we test the use of functional connectome fingerprinting as a way to select pre- and post-processing parameters. We hypothesized that whichever parameters yielded the best fingerprint accuracies would also improve the ICC across scans. Using five datasets from the Consortium for Replicability and Reproducibility, we found that higher identification accuracies were achieved when using: (I) global signal regression; (II) finer brain parcellations; (III) cortical regions compared to subcortical and cerebellar structures; (IV) medial frontal and frontoparietal networks relative to the whole-brain; (V) discriminative edges; (VI) longer scan duration; and (VII) lower sample size. We observed that the ICC was consistently “poor” across the five datasets even with the application of two optimal fingerprint-informed pipelines. The fingerprint-informed pipelines may enable comparison, benchmarking, and adjudication of functional connectivity-based analysis pipelines or novel analytic approaches, as a means to enhance their reproducibility in heterogeneous populations.

**Key Points:**

1. Connectome-based fingerprinting can provide a useful testbed for reproducible functional connectivity analysis pipelines.
2. Fingerprint-informed pipelines offer an intuitive and less resource intensive way to select data pre-/post-processing parameters for improving the reproducibility of the functional connectome.
3. Connectome-based fingerprinting offers an alternative approach to test-retest reliability.

## 1 Introduction

Despite the promise of increasingly representative and deeply phenotyped consortia, reproducible functional connectivity-based biomarkers have remained elusive. The identification of clinically useful brain-based biomarkers requires that our measures of the brain (e.g., functional connectivity) and of behavior (e.g., clinically important symptoms or behaviors) are reproducible. One plausible way of achieving brain-based biomarkers lies in employing precision fMRI strategies by “deep” sampling functional data across repeated measurements (e.g., Midnight Scan Club [MSC]; Gratton et al., 2020). This is because a key component of reproducibility is reliability (Castellanos et al., 2013) — the stability of a measure over time and under similar plausible scenarios (Mellinger and Hanson, 2020; Heale and Twycross, 2015). The reliability of a measure establishes an upper limit on its validity, where validity is measured in terms of its correlation with a target measure (e.g., behavior; Noble et al., 2019). Investigating the reliability of functional connectivity metrics is therefore crucial to advancing reproducible and clinically meaningful connectivity-based biomarkers.

One challenge to assessing functional connectivity metrics is the lack of a “ground truth” against which to evaluate their validity and which could therefore serve as a basis for comparing methods and pipelines for computing functional connectivity (Zuo et al., 2019). For example, while functional connectivity shows good correspondence with measures of anatomical or structural connectivity (e.g., as revealed by retrograde tracer studies in non-human animals, or diffusion-based connectivity studies in humans) (Litwińczuk et al., 2022; Margulies et al., 2009), it is clear that functional connectivity metrics reflect more than is captured by these anatomical measures, so correspondence with such measures alone cannot be used to evaluate functional connectivity metrics or methods for their computation. In the absence of a ground truth for evaluation of validity, many studies have focused on test-retest (TRT) reliability as an outcome measure; if a particular functional connectivity metric or analysis step or pipeline is advantageous, it should exhibit improved TRT reliability. The most consistently used index to evaluate TRT reliability of functional connectivity-based measures is the intraclass correlation coefficient (ICC; Noble et al., 2019; Shehzad et al., 2009; Shrout and Fleiss, 1979). While the ICC has been informative, it has a number of limitations. First, it is typically computed either as a voxelwise or an edge-level measure and exhibits significant variation both within and between networks (e.g., Shehzad et al., 2009); when an average value is computed across the brain, it is often quite low, suggesting that the TRT reliability for functional connectivity-based measures is poor-to-fair, at best (see Noble et al., 2019 for a review). This may, in part, reflect the fact that ICC is computed as the ratio of between-subject variance to within-subject variance (Shehzad et al., 2009). Since between-subject variance tends to account for majority of the variability in the ICC (Noble et al., 2019), achieving *high* between-subject and *low* within-subject variances for all functional connectivity measures remains a challenge (Tozzi et al., 2020; Noble et al., 2021; Chen et al., 2018). For these reasons, the ICC might not provide the optimal testbed for arbitrating between analysis pipelines.

Here, we test a straightforward alternative to ICC for comparing analyses and pipelines — connectome fingerprint identification. This process, first described by Finn et al. (2015), involves treating the functional connectivity profile (connectome) of an individual as a “fingerprint”. Fingerprint identification is performed by matching (using correlation) an individual’s connectome fingerprint derived from a given fMRI scan (taken from a “database” set of scans) with their connectome fingerprint derived from another, separate scan (taken from a “target” set of scans, typically acquired during a different scan session). If the correlation between a given individual’s two functional connectivity matrices is greater than the correlation between the individual’s database connectome and the connectome of other individuals in the target dataset, then it is considered a “match” — a correct identification (ID). If the self-self correlation is not greater than the self-other correlations, the individual has not been successfully identified by the matching algorithm. Studies applying the connectome fingerprinting approach across populations and scan conditions show that mean ID accuracy across a given sample can be >95% (St-Onge et al., 2023; Vanderwal et al., 2017; Finn et al., 2015), providing a different perspective on the reproducibility of functional connectivity metrics, relative to the ICC. Given its applicability, straightforward calculation, and straightforward interpretability, we suggest that connectome fingerprint ID offers a simple, intuitive, and less computationally demanding alternative to the ICC for the assessment of analysis pipelines and novel methodologies. We extend this logic to hypothesize that if a particular analysis step or pipeline is advantageous in ways that relate to reliability and/or capturing intersubject differences in functional connectivity, then it should improve fingerprint ID.

Several papers have already shown that ID accuracy is influenced by a number of processing choices. For example, Finn et al. (2015) reported lower ID accuracy when a cortical parcellation scheme was used to derive the functional connectivity profiles, relative to a whole-brain atlas. This may reflect the impact of differential parcel resolutions or spatial coverage (cortical vs. subcortical regions). Indeed, fingerprint ID accuracy exhibits variation across functional networks, with the highest ID accuracies being found in the medial frontal and frontoparietal networks (Vanderwal et al., 2021; Cai et al., 2021; Jalbrzikowski et al., 2019; Horien et al., 2019; Byrge et al., 2019; Vanderwal et al., 2017; Finn et al., 2015). Somewhat predictably, restricting analyses to a small subset of the most distinct connectome edges (e.g., < 1%) also improves ID accuracy, producing near-ceiling accuracy (Byrge et al., 2019). Other factors known to affect ID accuracy include scan duration (Vanderwal et al., 2021; Vanderwal et al., 2017; Finn et al., 2015), parcellation resolution (Vanderwal et al., 2017), and participant age (Horien et al., 2019; Kaufmann et al., 2017), with the youngest participants showing lowest ID, reflecting poor connectome distinctiveness, particularly between sessions (Dufford et al., 2021). Recent work also indicates that denoising strategies (e.g., global signal regression (GSR), temporal censoring, ICA-AROMA) improve ID accuracy in early childhood (Graff et al., 2022). Importantly, and contrary to the benefits commonly inherent to increasing sample size, increasing sample size has been shown to *decrease* ID accuracy, making it more difficult to identify an individual from a group (Li et al., 2021; Waller et al., 2017).

The present study has twin aims. First, we aim to show how fingerprint accuracy can serve as a robust, easily computed, intuitive, and sensitive index for comparing or benchmarking pipeline parameters. To achieve this aim, we expand on recent efforts to maximize functional connectome identification accuracy by examining the contributions of data pre- and post-processing factors, as well as acquisition parameters, using five independent datasets spanning childhood to adulthood, obtained from the Consortium for Replicability and Reproducibility (Zuo et al., 2014). By identifying several factors that maximize ID accuracy across the independent datasets, we delineate components of a pipeline that is optimized for the reliable measurement of functional connectivity at the individual level. The use of fingerprint accuracy as part of a workflow that seeks to maximize the identifiability of individual connectomes may also help improve the identification of robust and reproducible brain-behavior relationships.

## 2 Materials & Methods

### 2.1 Datasets

Five datasets from the Consortium for Replicability and Reproducibility (CoRR; http://fcon_1000.projects.nitrc.org/indi/CoRR/html/samples.html) were used: Beijing Normal University (BNU), New York University (NYUado, NYUadu), Southwest University (SWU), and University of Pittsburgh School of Medicine (UPSM). Ethical approval was obtained from the relevant committee at each imaging site. Each dataset included resting-state fMRI scans obtained from two imaging sessions either on the same day or several months or years apart. Demographics and fMRI acquisition parameters are provided in **Table 1**.

**Table 1.**
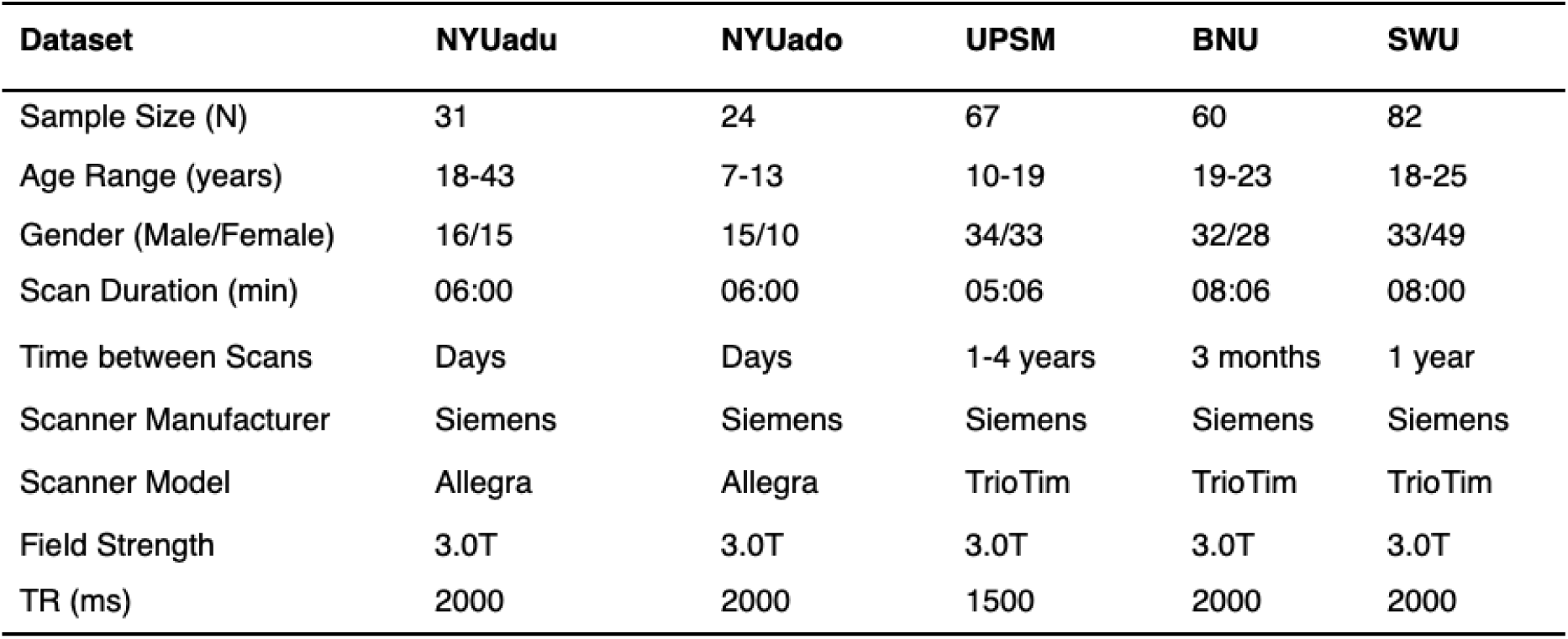
Demographic information and fMRI parameters for the five datasets obtained from CoRR that were used in this study. All sites contributed two resting state scans per participant.

### 2.2 Preprocessing

All data were preprocessed with the same basic pipeline (Yan et al., 2013a; Yan et al., 2013b; Craddock et al., 2013). Structural and functional data were preprocessed using AFNI (http://afni.nimh.nih.gov/afni/), FSL (http://fsl.fmrib.ox.ac.uk/), and ANTs (http://stnava.github.io/ANTs).

The preprocessing pipeline included: (1) volume-based motion correction; (2) grand mean scaling; (3) linear and quadratic detrending; (4) nuisance signal regression on 24 motion parameters (i.e., 3 translational, 3 rotational, their first derivatives, and the squares of each of these terms) and 12 nuisance regressors (i.e., white matter, CSF, global signal, though global signal was also a factor of interest so for some analyses, GSR was excluded from the nuisance regressors), their squares, and the squares of their first derivatives); (5) spatial smoothing using a Gaussian kernel at 6mm FWHM; and (6) temporal filtering (0.009 ≤ *f* ≤ 0.1 Hz). Functional to session-averaged anatomical co-registration was performed using boundary-based registration in FSL (Greve and Fischl, 2009). Diffeometric normalization of individual anatomical to the MNI152 standard space was carried out using ANTs; the same non-linear transformation was applied to the fMRI data. Participants were excluded if either scan exhibited root-mean-square framewise displacement (rmsFD) > 0.20 mm (Jenkinson et al., 2002).

### 2.3 Data Pre- and Post-Processing Parameters

To demonstrate how fingerprint ID accuracy can be used to test pipeline parameters, and to compare the utility of ID accuracy vs. ICC for this purpose, we examined how several data acquisition, pre-, and post-processing parameters affected ID accuray and ICC (**Figure 1**), specifically: (1) Global Signal Regression; (2) parcellation schemes (i.e., scheme, resolution, spatial coverage); (3) network organization; (4) use of discriminative edges; (5) scan duration; and (6) sample size.

**Figure 1.**
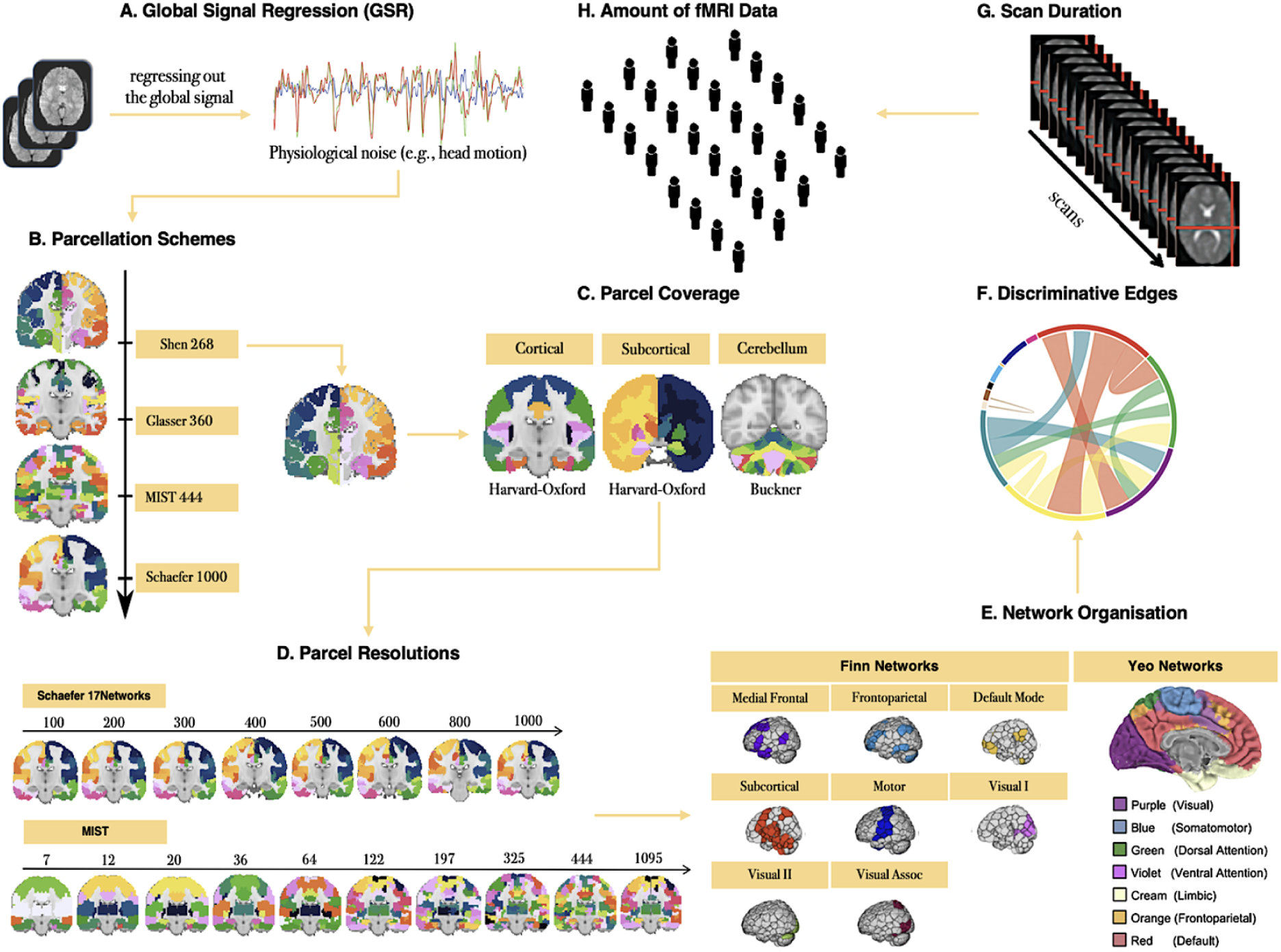
Analysis framework to delineate an optimal pipeline for maximizing connectome-based identification. A. Applying global signal regression (GSR) as a denoising strategy. B. Delineating the functional connectome with four publicly available parcellation schemes such as Shen 268, Glasser 360, MIST 444 and Schaefer 1000-17Networks. C. Delineating cortical and subcortical regions (Harvard-Oxford atlas and Buckner atlases). D. Computing the functional connectome using multi-resolution parcellation schemes with 8-10 levels of dimensionality. E. Examining ID within “Finn” Networks and “Yeo” Networks. F. Using the top 1-75% of most discriminative edges. G. Examining scan duration — from 0.5 min to the full acquisition time of each dataset. H. Examining the impact of sample size (*N*) by pooling data across the five datasets and resampling such that *N* ∈ {20, 40, 60, 80, 100, 120, 140, 160, 180, 200, 220, 240, 264}.

#### 2.3.1 GSR

Although debated, GSR is a highly effective denoising strategy (Parkes et al., 2018; Satterthwaite et al., 2017; Ciric et al., 2017). It remains the most effective strategy for reducing the associations between motion and functional connectivity-based measures (Power et al., 2017; Power et al. 2014; Yan et al., 2013a; Yan et al., 2013b) in addition to increasing the biological plausibility of patterns of functional connectivity (e.g., Weissenbacher et al., 2009; Fox et al., 2009). We examined the impact of including vs. excluding the global signal from the 36-parameter nuisance signal regression step during preprocessing.

#### 2.3.2 Parcellation Schemes (Selection, Spatial Coverage & Resolution)

Parcellation schemes differ in aspects such as spatial coverage (e.g., cortex, subcortex), space (e.g., volume, surface), and resolution (e.g., 10-1000). We parcellated the fMRI data using four publicly available brain atlases: Shen 268 (Shen et al., 2013), Glasser 360 (Glasser et al., 2016), MIST 444 (Urchs et al., 2017), and Schaefer 1000-17Networks (Schaefer et al., 2018), to examine whether these different commonly applied schemes are associated with differences in ID accuracy. We also compared ID accuracies with cortical-only parcellations to parcellations that included subcortical and cerebellar regions. Finally, we assessed the relationship between resolution and ID accuracy by examining two parcellation schemes with 8 (Schaefer 17Networks, 100-1000) and 10 (MIST, 7-1095) levels of dimensionality.

#### 2.3.3 Network Organization

Several studies have demonstrated that some functional networks have stronger predictive power in facilitating the identification of an individual (Jalbrzikowski et al., 2019; Horien et al., 2019; Finn et al., 2015). To assess differences between functional networks in terms of ID accuracy, we examined 8 functional “Finn” Networks: (1) medial frontal; (2) frontoparietal; (3) default mode; (4) subcortical and cerebellum; (5) motor; (6) visual I; (7) visual II; and (8) visual association (Finn et al., 2015) and 7 “Yeo” Networks: (1) ventral attention; (2) dorsal attention; (3) frontoparietal; (4) default mode; (5) somatomotor; (6) limbic; and (7) visual (Yeo et al., 2011).

#### 2.3.4 Discriminative Edges

Fingerprint identification is influenced by the degree to which different edges contribute to the identification of an individual (Horien et al., 2019; Byrge et al., 2019; Vanderwal et al., 2017; Finn et al., 2015). We employed the discriminative power (DP) measure to quantify the usefulness of an edge in successfully identifying an individual from a group. The DP of a given edge reflects the empirical probability of an individual being more correlated to themself relative to another individual based on the same edge. It is computed as the product of the edge values between sessions from an individual and compared with the product of the edge values between sessions from other individuals in a dataset (see Horien et al., 2019, Finn et al., 2015 for details). An edge with a high DP means that the within-subject product is greater than the between-subject product across all the participants and that it is useful for the identification of an individual. To compute ID accuracy with the most discriminative edges across all individuals in the five datasets, we selected the edges whose DP values fell between the 25^th^ and 99^th^ percentiles of the functional connectome, i.e., we retained the top 1-75% of edges in the whole-brain connectome to compute ID accuracy for each dataset.

#### 2.3.5 Scan Duration

Converging evidence has revealed that the reliability of functional connectivity-based measures depends on scan duration (Noble et al., 2017; Mueller et al., 2015; Birn et al., 2013; Anderson et al., 2011; Van Dijk et al., 2010) and that longer scan duration improves ID accuracy (Vanderwal et al., 2021; Vanderwal et al., 2017; Airan et al., 2016; Finn et al., 2015). We assessed the relationship between scan duration and ID accuracy by increasing the scan lengths from 0.5 min to the full acquisition time of each dataset (scan parameters for each site in **Table 1**). Of note, the range of the scan lengths is limited based on the relatively short scan durations of the five datasets (e.g., the maximum scan length is approximately 8 min derived from the BNU and SWU datasets).

#### 2.3.6 Sample Size

While sample size typically improves the reliability of functional connectivity-based measures (Cho et al., 2020; Elliott et al., 2019), identification accuracy tends to decrease with increasing sample sizes (Li et al., 2021b; Waller et al., 2017). To assess the impact of sample size on ID accuracy, and to avoid site/sample effects, we pooled the five datasets (*N* = 264), then randomly selected subsets of individuals such that *N* ∈ {20, 40, 60, 80, 100, 120, 140, 160, 180, 200, 220, 240, 264} *without* replacement and computed ID accuracy over 100 of each *N* iterations.

### 2.4 Functional Connectivity Matrices **—** Fingerprints

For every individual and each scan, the mean timeseries of each region of interest (ROI) within a given parcellation scheme was extracted and the Pearson’s correlation coefficient (*r*) between all possible ROI pairs was computed, yielding two functional connectivity matrices (fingerprints) per individual that were subsequently used in the ID procedure.

### 2.5 Identification Procedure

To identify an individual based on their functional connectivity fingerprints, we followed the identification procedure as previously described (Finn et al., 2015). Identification was performed for each dataset separately by creating a “database” **D** = {*X*, *i* = 1, 2, 3….*N*} where *X_i_* refers to the vectorised functional connectivity matrix for Session 1 and the subscript *i* represents the *i*^th^ participant in **D**. Iteratively, *X_i_* is compared with functional connectivity matrix *X_j_* { *j* = 1, 2, 3….*N*} from the Session 2 target set, **T**, using correlation. Correlations were rank ordered, and an individual was assigned a binary identification (BID) score of 1 if the predicted identity *X_j_* (rank = 1) in **T** matched the true identity *X_i_* of the individual in **D**, otherwise the BID score was 0. Mean BID accuracy for each dataset and each combination of analysis factors was computed as the number of correctly identified participants divided by *N* (sample size). To reach a final ID accuracy per dataset, we recomputed mean BID by exchanging the roles of the database-target matrix (**D** ↔ **T**), and averaging across the two mean BIDs.

In addition to BID, we computed the relative rank (RR), which is a continuous measure ranging from 0 to 1 that quantifies the degree of “confusion” for inaccurately identified individuals. RR was calculated as the *j*^th^ position of the rank ordered correlation that matches the predicted identity *X_j_* in **T** with *X_i_* in **D** divided by *N*. For example, if the ranked ordered correlation of participant 1 in the SWU dataset is at the first position, then RR = 1/82 = 0.012 whereas if the ranked ordered correlation for the same participant is at the last position, then RR = 82/82 = 1. The fewer individuals inaccurately ranked above their true identity, the lower the degree of confusion, and lower the RR (RR → 0).

### 2.6 TRT Reliability

We computed TRT reliability, as indexed by ICC, for each edge and each dataset. We applied the ICC (2,1) model to refer to “absolute agreement” for which the sources of error are known (e.g., multiple sessions) and modeled as random effects (Noble et al., 2019). For each dataset, the mean ICC was obtained by averaging the edgewise ICC values in the functional connectome in addition to setting the negative values to 0 to indicate complete unreliability (Noble et al., 2019; Noble et al., 2017; Bartko, 1976).

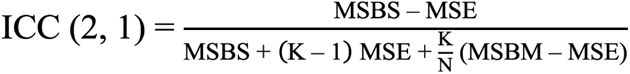

where MSBS = mean square between subjects, MSBM = mean square between measurements, and MSE = mean square error (Liljequist et al., 2019; Koo and Li, 2016):

### 2.7 Comparing Analysis Pipelines Using Fingerprint ID vs. ICC

For each dataset, the identification accuracy was computed with each of the data acquisition, pre-processing, and post-processing parameters, and then compared with a “standard” functional connectivity pipeline as described in Sections 2.2 and 2.4. The standard pipeline comprised a 36-parameter confound regression model including GSR and its derivatives, whole-brain Shen 268 parcellation, full scan length, and full sample size of each dataset. On the basis of the *highest* contributions of the parameters on ID accuracy (i.e., BID → 100% and RR → 0%), we identified two optimal fingerprint-informed pipelines which are as follows:

- <u>Pipeline 1</u>: minimal pre-processing, GSR, Schaefer 1000-17Network parcellation, full scan length, full sample size;
- <u>Pipeline 2</u>: minimal pre-processing, GSR, frontoparietal networks (medial frontal + frontoparietal) derived from “Finn” Networks, full scan length, full sample size.

For each dataset, the mean ICC was further computed with Pipeline 1 and Pipeline 2. To demonstrate the utility of the two fingerprint-informed pipelines, they were employed to derive the functional connectivity matrices of the participants for their two fMRI scans for each dataset. Given that we assessed how fingerprint ID accuracy can be applied to test analysis pipeline parameters, we did not perform permutation testing to assess the statistical significance of the ID accuracies themselves across the five datasets.

### 2.8 Data Sharing and Code Availability

The analyses in this study were conducted in Python and the codes are available via our GitHub repository (https://github.com/JRam02/conn_fingerprint_neurodev). We used publicly available packages such as pingouin (https://pingouin-stats.org/api.html), scikit-learn (https://scikit-learn.org/stable/), and SciPy (https://www.scipy.org). We also used Python libraries for data manipulation and visualization such as Numpy (https://numpy.org), Pandas (https://pandas.pydata.org), Matplotlib (https://matplotlib.org), and Seaborn (https://seaborn.pydata.org). The ICC analysis was conducted using the intraclass_corr package in pingouin. The fMRI data of the five datasets that were used in this study are available on the CoRR platform. We also provide the mean timeseries of the participants for each dataset via our GitHub repository.

## 3 Results

### 3.1 Identifiability of the individual functional connectome

Using whole-brain functional connectomes collected from two scans per participant derived from CoRR, we initially applied a standard functional connectivity pipeline which involves minimal data pre-processing (including GSR), post-processing (i.e., whole-brain Shen 268 parcellation), and acquisition (i.e., full scan duration and sample size) parameters to compute the identification accuracy across the five datasets. We first computed binary identification (BID) such that the BID accuracy is 1 if an individual is successfully identified and 0 if they are not, using all five datasets. BID accuracies ranged from 42% to 77% which were averaged across the two database-target matrix configurations (**Figure 2A** and **Figure 2B**). On average for both directions (**D** → **T** and **T** → **D**), the NYU developmental dataset (NYUado; *N* = 24) achieved the strongest identification accuracy (BID = 77%) whereas the UPSM adolescent group (*N* = 67) had the lowest (BID = 42%). We then employed relative rank (RR) to measure the degree of confusion for misidentifying an individual from a group such that the lower the RR, the lower the confusion for incorrectly identifying that individual. RR showed that the degree of confusion varied from 6% to 13% for the first direction (**D** → **T**) (**Figure 2A**), and 4% to 16% for the second direction (**T** → **D**) (**Figure 2B**). On average, the adult dataset (SWU) achieved the lowest degree of confusion for incorrectly identifying an individual (RR = 5%) between sessions whereas the UPSM adolescent dataset exhibited the highest degree of confusion (RR = 15%).

**Figure 2.**
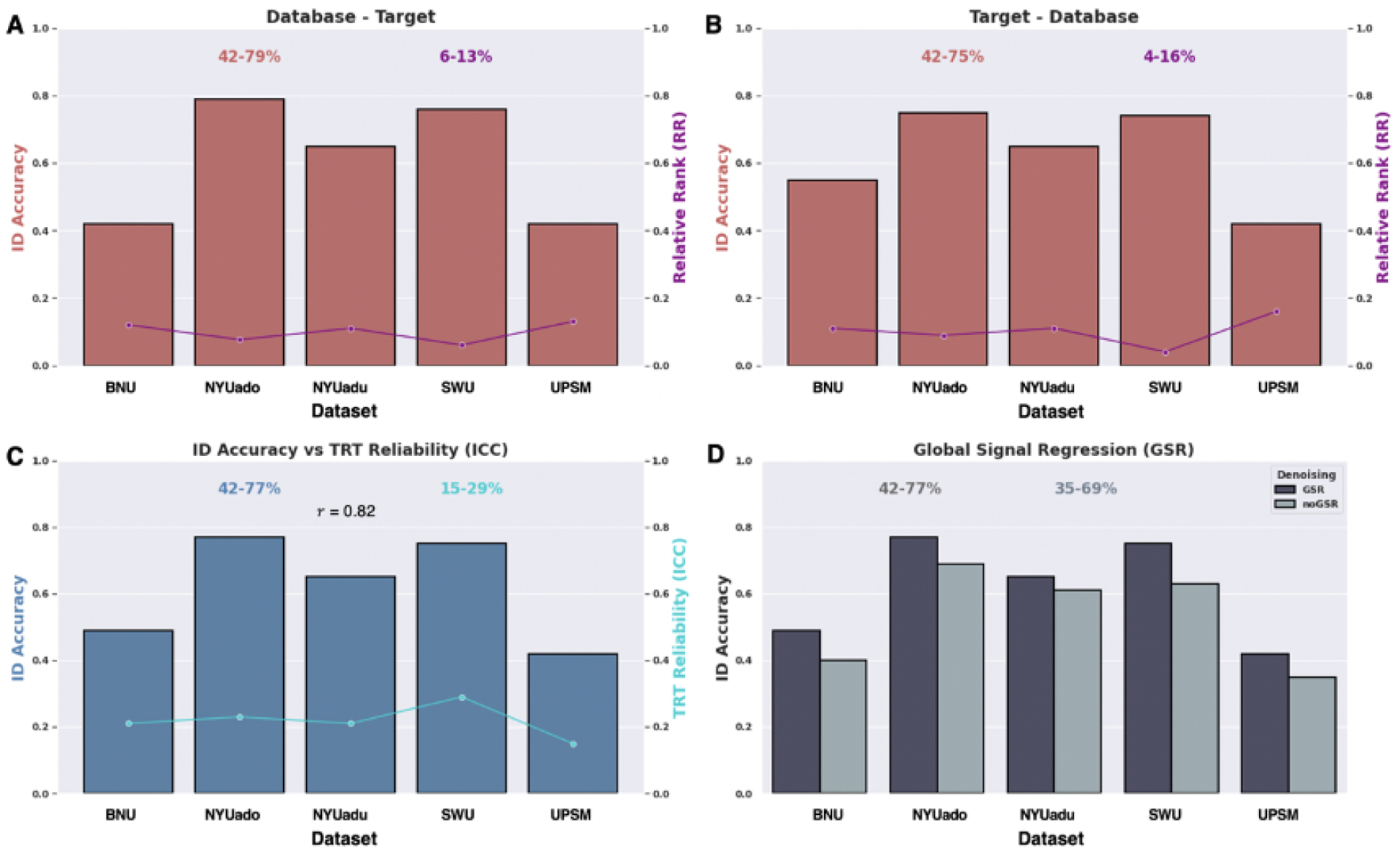
(A) ID accuracy derived from BID in addition to the degree of confusion as indexed by RR for the database-target matrix configuration. (B) ID accuracy derived from BID in addition to the degree of confusion as indexed by RR for target matrix-database configuration. (C) Reliability estimates derived from the fingerprint approach using BID and the ICC(2,1) model. The Pearson’s correlation coefficient (*r*) between ID accuracy and ICC is 0.82. (D). ID accuracy derived from BID with and without applying GSR as a denoising strategy. The five independent datasets are as follows: BNU (*N* = 60), NYUado (*N* = 24), NYUadu (*N* = 31), SWU (*N* = 82), and UPSM (*N* = 67).

### 3.2 TRT reliability of the functional connectome is consistently poor

We then evaluated the TRT reliability of the functional connectome by applying the ICC(2,1) model using the standard functional connectivity pipeline across the five datasets. We computed the average ICC across 268 x 268 edges for the Shen ROIs. We found that the mean TRT reliability was consistently “poor” (i.e., ICC < 0.4) across the independent datasets (**Figure 2.2C**). The highest mean TRT reliability was observed for the adult dataset (SWU; ICC = 0.29) whereas the lowest mean TRT reliability was found for the adolescent group (UPSM; ICC = 0.15). Although the ICC and BID accuracy can only be compared qualitatively, the two metrics track each other well (*r* = 0.82; **Figure 2.2C**).

### 3.3 Applying GSR as a denoising strategy improves identification accuracy

While investigating the impact of data pre-processing factors on identification accuracy across the five datasets, we assessed the contribution of GSR by computing identification accuracy following pre-processing *with* and *without* GSR (**Figure 2D**). For the purpose of this comparison, all other noise regressors were held constant (i.e., the global signal was the only regressor removed from the 36-parameter nuisance signal regression). The range of the BID accuracies was higher *with* GSR (42-77%) than *without* GSR (35-69%). GSR significantly improved the BID accuracies across all five datasets (paired sample t-test; t (4) = 6.14, *P* = 0.004).

### 3.4 Finer cortical parcellations yield stronger identification accuracy

To understand how different parcellation features influence the identification accuracy, we computed functional connectivity using four parcellation schemes with different spatial resolutions and spatial coverage. We found that the BID accuracies differed significantly across parcellation schemes (repeated measure one-way ANOVA test; F (3,12) = 31.1, *P* < 0.001). There was a tendency to achieve greater BID accuracies across the five datasets when the functional connectome was derived from finer parcellation schemes, i.e., parcellations with higher resolutions. While the lowest BID accuracies were obtained with the Shen 268 parcellation (42-77%), the highest BID accuracies were found with the Schaefer 1000-17Network parcellation (68-94%) (**Figure 3A**). Both Glasser 360 and MIST 444 parcellation schemes produced intermediate BID accuracies ranging from 55% to 86% across the five datasets (**Figure 3A**).

**Figure 3.**
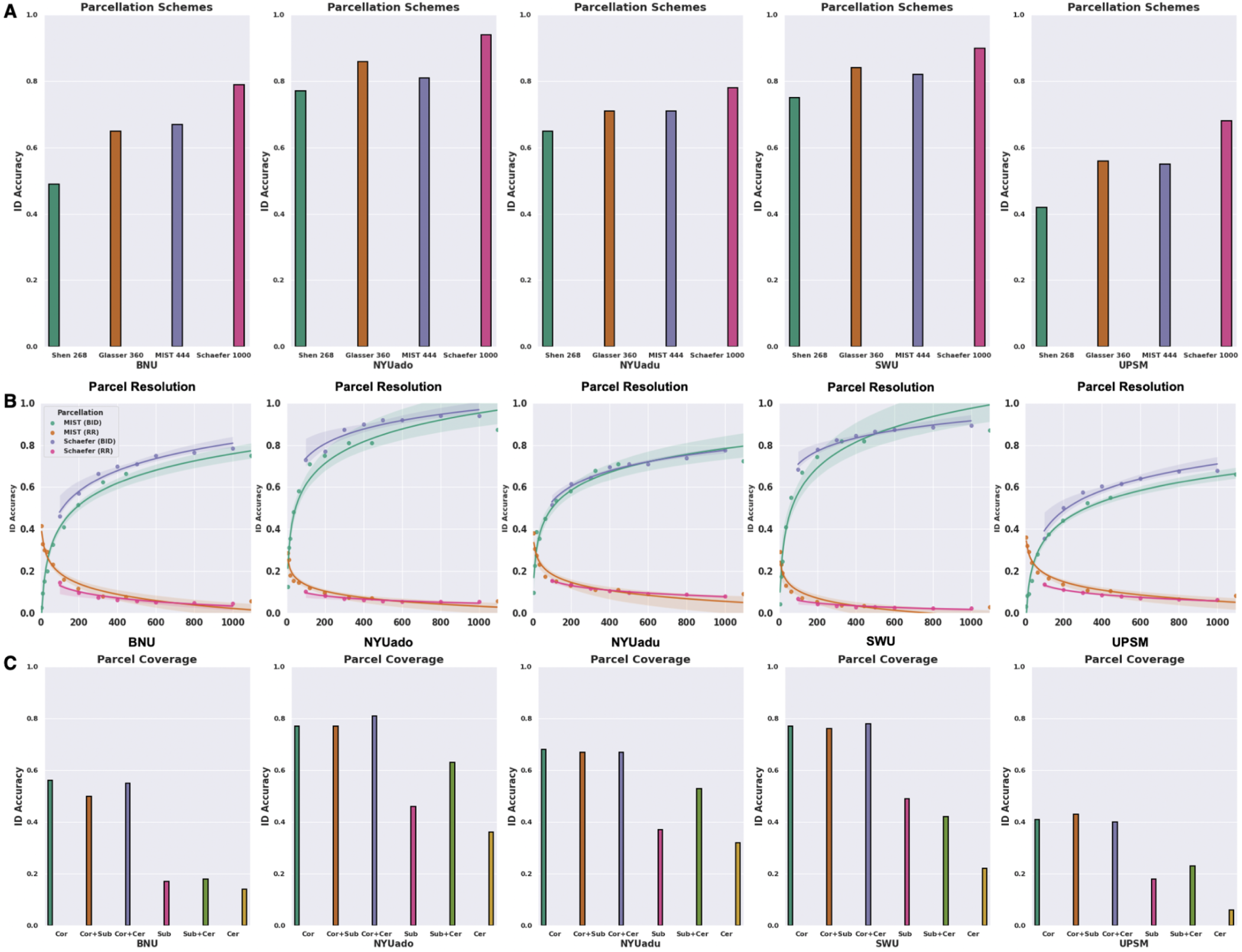
(A) ID accuracy obtained from BID across the five independent datasets using four publicly available parcellation schemes including Shen 268, Glasser 360, MIST 444 and Schaefer 1000-17Networks. (B) Relationship between parcel resolution and ID accuracy measured by BID in addition to the degree of confusion as indexed by RR using 8 and 10 levels of parcel granularity derived from the Schaefer 17Networks and MIST parcellation schemes across the five independent datasets. (C) ID accuracy derived from BID based on the spatial coverage of the functional connectome covering cortical, subcortical and cerebellar structures across the five independent datasets. Cor: cortical; Cor+Sub: cortical and subcortical; Cor+Cer: cortical and cerebellum; Sub: subcortical; Sub+Cer: subcortical and cerebellum; and Cer: cerebellum.

To assess the effect of resolution more cleanly and within the same parcellation scheme, we applied the multi-resolution Schaefer 17Networks (i.e., 100-1000) and MIST (i.e., 7-1095) parcellations to examine the relationship between parcel resolution and identification accuracy. There was a non-linear relationship between parcel resolution and BID accuracy that was evident across the five datasets (**Figure 3B**). BID showed that there were large gains in ID accuracy as parcel resolution increased from 7 to 300, with smaller but still significant gains thereafter (**Figure 3B**). Similarly, there was a non-linear relationship between parcel resolution and RR across the five datasets such that there was a rapid decline in RR from 7 to 300 parcels, and then plateaued until reaching the finest resolutions (**Figure 3B**).

We also investigated how using regions from the cortex, subcortex, and cerebellum influences identification accuracy. Fingerprint BID accuracies differed significantly across different combinations of the spatial coverage (repeated measure one-way ANOVA test; F (5,20) = 54.4, *P* < 0.001). The highest BID accuracies were derived when the functional connectome included cortical regions (41-77%), with the addition of subcortical and cerebellar regions making little-to-no difference to BID accuracy (**Figure 3C**). Fingerprints computed on the basis of subcortical (17-49%) and/or cerebellar (6-36%) structures alone were associated with much lower BID accuracies across the five datasets (**Figure 3C**). The higher fingerprint BID accuracies associated with the cortical regions could be attributed to their macroscale divisions having larger number of edges, as we found significant associations between number of edges and BID accuracies for 4 out of 5 datasets (except for the BNU dataset).

### 3.5 Identification accuracy across functional brain networks

To examine variation in fingerprint identifiability across functional networks, we compared the BID accuracies across eight “Finn” networks and seven “Yeo” networks. For the “Finn” networks, BID accuracies were highest when the fingerprints were restricted to the medial frontal (except for the UPSM dataset) and frontoparietal networks, relative to the other functional networks (**Figure 4A**). In contrast, fingerprinting using the visual networks (i.e., visual I, visual II, visual association) produced the lowest BID accuracies (**Figure 4A**). Given previous research showing that combining medial frontal and frontoparietal networks improved BID accuracies (Horien et al., 2019; Finn et al., 2015), we examined the impact of combining these functional networks into the “frontoparietal networks”. Restricting the fingerprint to these frontoparietal networks was associated with significantly greater identification accuracies compared to the whole-brain functional connectome (paired sample t-test; t(4) = 2.83, *P* = 0.047). When we applied the “Yeo” networks, BID accuracies revealed that the fingerprints restricted to the frontoparietal (except for the NYUadu dataset; 57-87%) and default mode networks outperformed the other functional networks across the five datasets (**Figure 4B**). Across the five datasets, fingerprinting with the limbic network exhibited the lowest BID accuracies (21%-67%) followed by fingerprinting with the visual network (25%-69%) (**Figure 4B**). When testing if the advantages in fingerprint BID accuracies for the frontoparietal networks might be due to a network size (i.e., networks having larger number of edges), we found that for the “Finn” networks, only the SWU dataset produced a significant relationship between network size and BID accuracies (*r*_s_ = 0.72, *P* = 0.028). For the “Yeo” networks, only the NYUado (*r* = 0.79, *P* = 0.035) and UPSM (*r* = 0.77, *P* = 0.041) datasets yielded a significant relationship between network size and BID accuracies.

**Figure 4.**
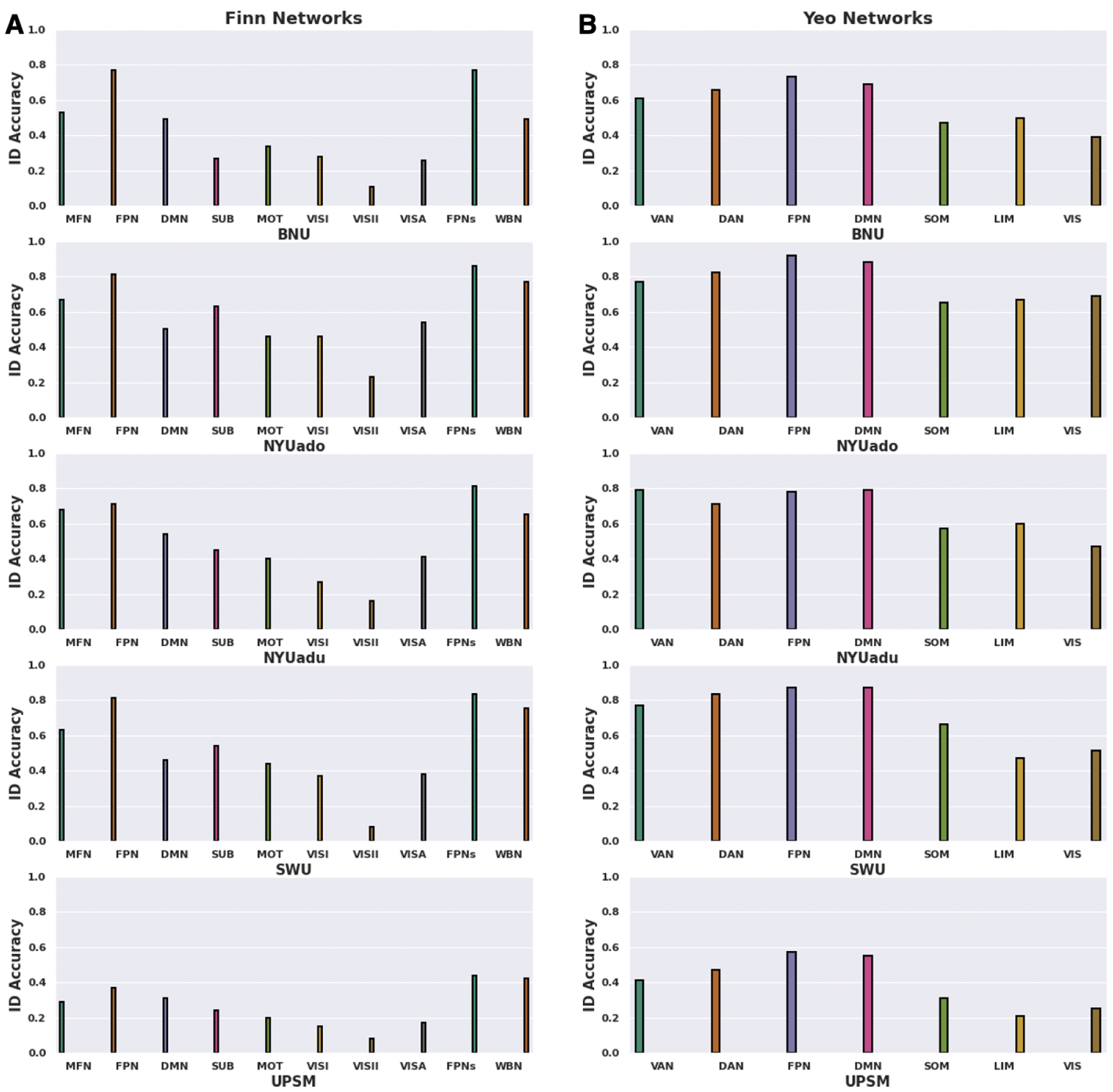
(A) ID accuracy derived from BID for the Finn Networks in addition to the combined medial frontal and frontoparietal networks [FPNs] and whole-brain connectome [WBN] across the five datasets. The Finn Networks are as follows: (1) medial frontal [MFN]; (2) frontoparietal [FPN]; (3) default mode [DMN]; (4) subcortical [SUB]; (5) motor [MOT]; (6) visual I [VISI]; (7) visual II [VISII]; and (8) visual association [VISA]. (B) ID accuracy derived from BID for the Yeo Networks across the five datasets. The Yeo Networks are as follows: (1) ventral attention [VAN]; (2) dorsal attention [DAN]; (3) frontoparietal [FPN]; (4) default mode [DMN]; (5) somatomotor [SOM]; (6) limbic [LIM]; and (7) visual [VIS]

### 3.6 Greater number of discriminative edges yields stronger identification accuracy

To examine whether restricting the fingerprint to discriminative edges improves identification accuracy, we restricted the fingerprint to those edges whose DP values fell between the 25^th^ and 99^th^ percentiles of the functional connectome for each dataset (i.e., selecting the top 1-75% of all edges, ordered by discriminality). We observed a non-linear relationship between the number of discriminative edges and BID accuracies across the five datasets (**Figure 5**). When the fingerprint was restricted to the top 1% of DP edges, BID accuracies (21-64%) were significantly lower compared to using all the edges from the whole-brain connectome (paired sample t-test; t (4) = 7.21, *P* = 0.002). When the fingerprint was computed with the top 10% of DP edges, BID accuracies (37-77%) were significantly higher compared to the top 1% of DP edges (paired sample t-test; t (4) = 8.22, *P* = 0.001). On the other hand, when the fingerprint was computed with the top 75% of DP edges, BID accuracies slightly improved (43-77%) compared to the whole-brain connectome. However, this improvement was not significantly different from the whole-brain connectome (paired sample t-test; t (4) = 1.84 *P* = 0.14).

**Figure 5.**
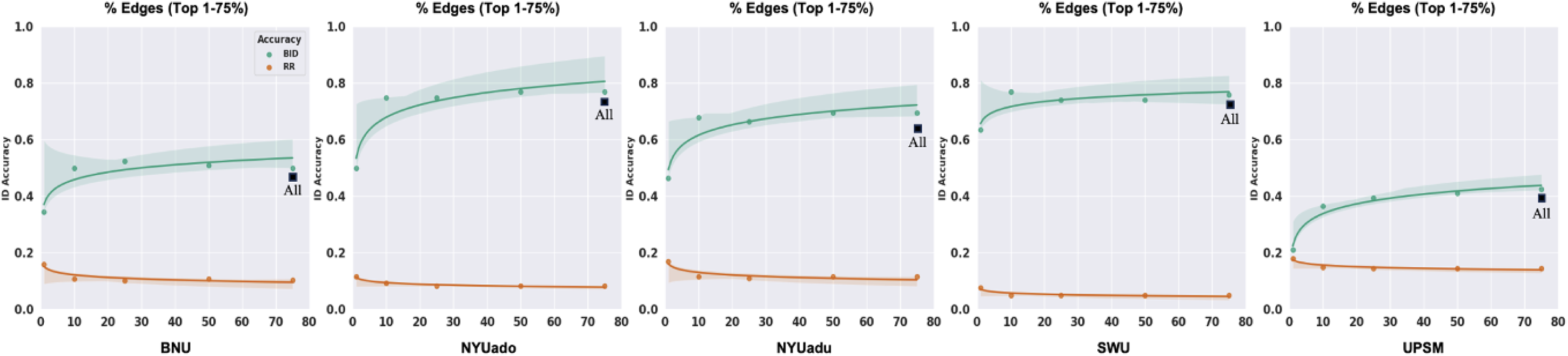
Relationship between number of discriminative edges and ID accuracy derived from BID in addition to the degree of confusion as indexed by RR across the five independent datasets. The discriminative edges are selected based on their DP values falling between the 25^th^ and 99^th^ percentiles of the whole-brain connectome, i.e., the top 1-75% of edges in the whole-brain connectome were retained to compute the ID accuracy of each dataset. The black box represents the ID accuracy derived from BID when all the edges in the whole-brain connectome were used for each dataset.

Similar observation was noted when comparing the BID accuracies from the top 10% of DP edges to the whole-brain connectome (paired sample t-test; t (4) = -0.14, *P* = 0.90). The non-linear relationship between number of discriminative edges and BID accuracy is striking — there is a large jump in BID accuracies between 1% and 10%, but little-to-no improvement after that, suggesting that good BID accuracies can be achieved with as little as the top 10% of the most discriminative edges. Of note, the potential benefits of using the most discriminative edges to perform fingerprinting can be outweighed by the computational challenge in estimating the DP for high-resolution parcellation schemes. For example, a coarser parcellation scheme such as the Shen 268 has 35,778 edges whereas a finer parcellation scheme like the Schaefer 1000-17Networks has 499,500 edges. Given that DP is computed as the product of an edge value from Scan 1 and Scan 2 from the same individual and compared with the product of the same edge value from Scan 1 and Scan 2 from other individuals, identifying the most discriminative edges based on their DP values will be feasible for smaller parcellation sets (e.g., 35,778 edges) but will create computational difficulties at larger (e.g., 499,500 edges).

### 3.7 Longer scan duration facilitates higher identification accuracy

We evaluated the effort of acquiring increasing scan length to boost identification accuracy by varying the scan duration from 0.5 min to the full acquisition length of each dataset. There was a non-linear relationship between scan duration and identification accuracy across the five datasets (**Figure 6A**), such that BID accuracies were very negatively impacted by very short fMRI scans (e.g., 0.5 min — ranging from 5% to 21% across the five datasets). BID accuracies were optimal (42-77%) when the full acquisition length of each dataset was used to compute the functional connectivity matrices of each individual. RR revealed a similar pattern such that the degree of confusion for unsuccessfully identifying an individual was highest (23-41%) with short acquisition time (e.g., 0.5 min) but minimal (5-15%) when the entire scan length was used to derive the functional connectivity profiles of the individuals across the five datasets.

**Figure 6.**
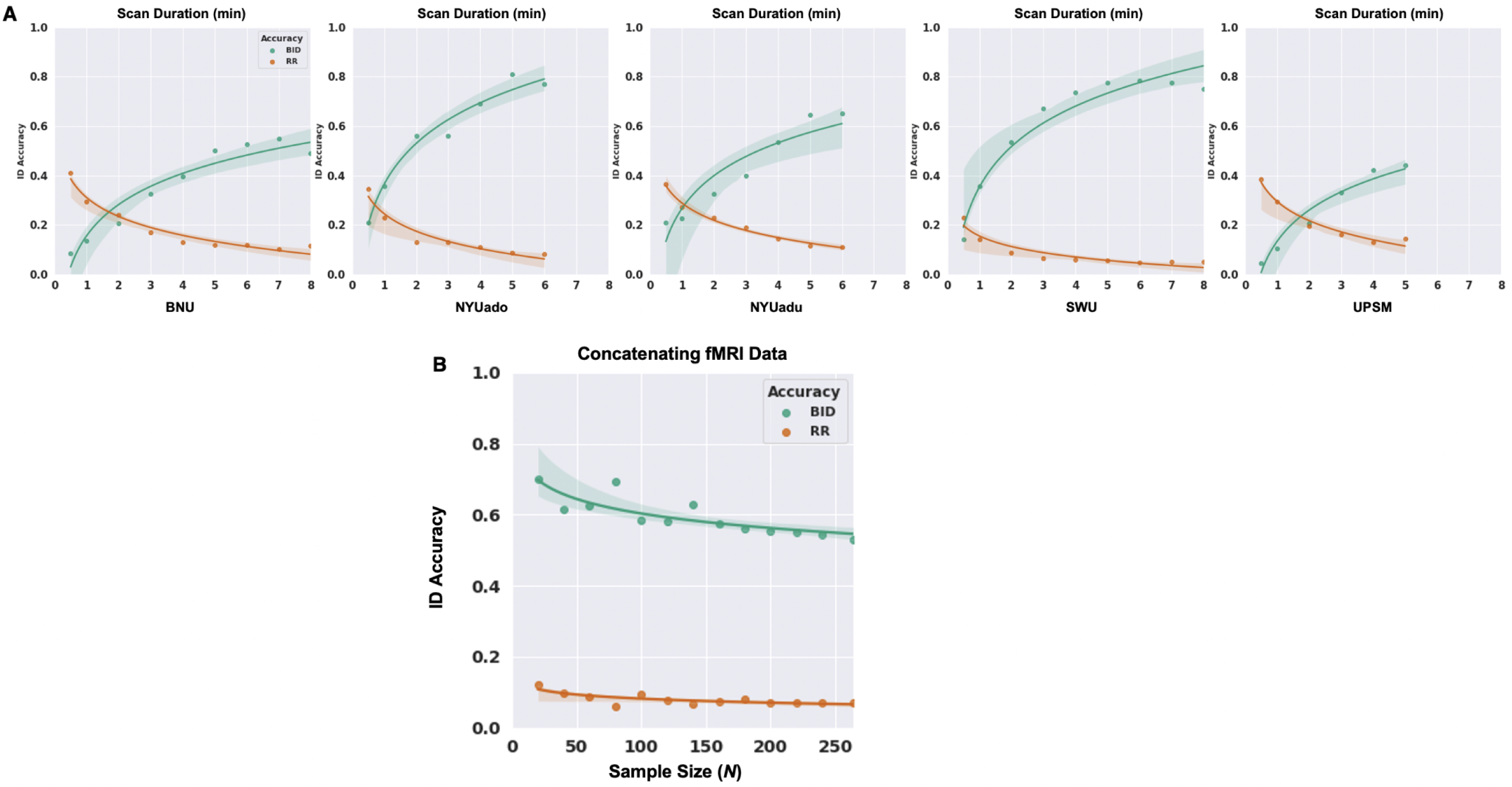
(A) Relationship between scan duration and ID accuracy measured by BID in addition to the degree of confusion as indexed by RR across the five independent datasets. The scan duration ranges from 0.5 min to the full acquisition time of each dataset. (B) Relationship between amount of resting-state fMRI data and ID accuracy measured by BID in addition to the degree of confusion as indexed by RR. The resting-state fMRI scans were concatenated across the five datasets (*N* = 264) for each imaging session, and then subsets of individuals were randomly selected across 100 iterations such that *N* ∈ {20, 40, 60, 80, 100, 120, 140, 160, 180, 200, 220, 240, 264}.

### 3.8 Increasing sample size does not necessarily improve identification accuracy

We also examined the effect of sample size (*N*) on identification accuracy by creating random subgroups of individuals *N* ∈ {20, 40, 60, 80, 100, 120, 140, 160, 180, 200, 220, 240, 264} across the five datasets. For each *N*, 100 samples were selected, and the average BID accuracy was computed across samples. There was a non-linear relationship between sample size and BID accuracy such that the ability to identify an individual based on their functional connectivity profiles decreased non-linearly as the sample size increased (**Figure 6B**), with the steepest decrease observed as sample size increased from 20 to 100, and a more gradual decrease in BID accuracy as sample size increased further to 264. The overall change across sample sizes was relatively small, however — BID accuracies ranged from 70% when *N* = 20 to 53% when *N* = 264. As a mathematical reality, increasing sample sizes also increases the potential for “confusion” (high correlation) amongst individuals — it increases the pool of potential matches, thus decreasing identification accuracy. The aggregated/subsampling approach produced higher BID accuracies for sample sizes equivalent to those of the UPSM, BNU, and NYUadu sites, relative to the BID accuracies for those sites alone, suggesting some site or sample effects that were overcome by the aggregating and subsampling approach. Further, and contrary to observations for the preceding manipulations, for which the trend for RR was inverted relative to the trend for the BID accuracy, RR *decreased* slightly but non-linearly from 12% to 7% as sample size increased (**Figure 6B**).

### 3.8 Selecting a fingerprint-informed pipeline

On the basis of the results presented above, we identified two optimal fingerprint-informed pipelines featuring those parameters associated with the *highest* BID accuracy (**Table 2**; **Figure 7**). The two pipelines have been described given that predictive networks such as the frontoparietal networks and finer cortical parcellation schemes such as the Schaefer 1000-17Network scheme, separately, encapsulate the most distinct features of an individual’s functional connectome. As expected, relative to the standard pipeline (minimal pre-processing, GSR, whole-brain Shen 268 parcellation, full scan length, and full sample size), BID accuracy was significantly higher for both Pipeline 1 (paired sample t-test; t (4) = 6.11, *P* = 0.004) and Pipeline 2 (paired sample t-test; t (4) = 2.83, *P* = 0.047). Notably, some datasets (e.g., NYUado; SWU) exhibited BID accuracies of 90% or above following processing with Pipeline 1.

**Figure 7.**
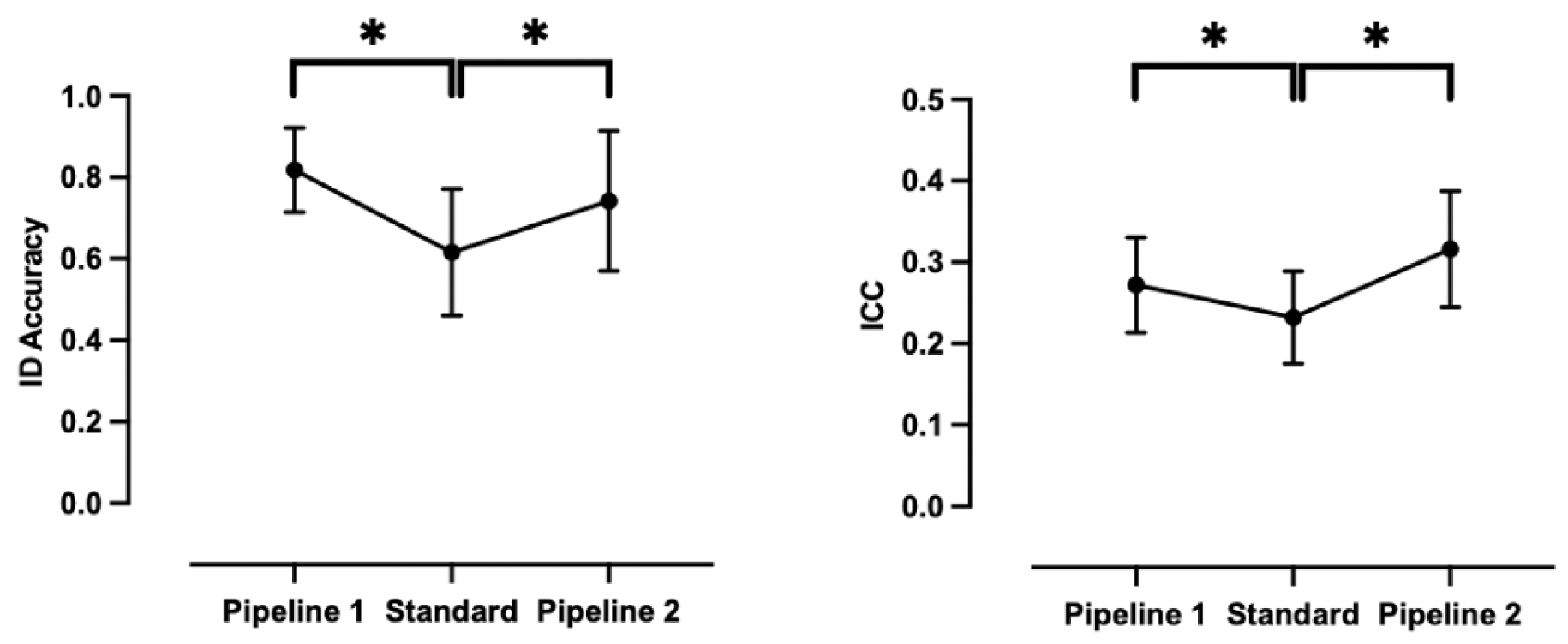
ID accuracy and TRT reliability of the functional connectome measured by BID and ICC, respectively, using the standard and optimal fingerprint-informed pipelines. Dots and error bars represent the mean and SD of the BID accuracies and ICC values. <u>Standard</u>: GSR + Shen 268 parcellation + full scan length + full sample size. <u>Pipeline 1</u>: GSR + Schaefer 1000-17Network parcellation + full scan length + full sample size. <u>Pipeline 2</u>: GSR + frontoparietal networks derived from Finn Networks + full scan length + full sample size. \**P* < 0.05, two tailed paired sample t-test. For ID accuracy, the effect sizes measured by Cohen’s d are: 1.5 (Pipeline 1) and 0.8 (Pipeline 2). For TRT reliability, the effect sizes measured by Cohen’s d are : 1.0 (Pipeline 1) and 1.6 (Pipeline 2).

**Table 2.** Gains in ICC and ID accuracy derived from BID across the five datasets derived from the standard and optimal fingerprint-informed pipelines. ⋄Standard: GSR + Shen 268 parcellation + full scan length + full sample size. *Pipeline 1: GSR + Schaefer 1000-17Network parcellation + full scan length + full sample size. ^‡^Pipeline 2: GSR + frontoparietal networks derived from Finn Networks + full scan length + full sample size.

| Pipeline | NYUadu | NYUado | UPSM | BNU | SWU |
| --- | --- | --- | --- | --- | --- |
| ◊Standard ICC | 0.21 | 0.23 | 0.15 | 0.21 | 0.29 |
| *Pipeline 1 ICC | 0.25 | 0.28 | 0.19 | 0.29 | 0.35 |
| ‡Pipeline 2 ICC | 0.29 | 0.33 | 0.21 | 0.35 | 0.40 |
| ◊Standard BID | 0.65 | 0.77 | 0.42 | 0.49 | 0.75 |
| *Pipeline 1 BID | 0.78 | 0.94 | 0.68 | 0.79 | 0.90 |
| ‡Pipeline 2 BID | 0.81 | 0.86 | 0.44 | 0.77 | 0.83 |

### 3.9 Applying functional connectome fingerprinting informed pipelines to improve ICC

To further assess the utility of these two fingerprint-informed pipelines, we computed TRT reliability as indexed by the ICC for each dataset following the application of these pipelines to compute functional connectivity (**Table 2**; **Figure 7**). The mean ICCs increased significantly with Pipeline 1 (paired sample t-test; t (4) = 7.22, *P* = 0.002) and Pipeline 2 (paired sample t-test; t (4) = 7.22, *P* = 0.002), although it is worth nothing that the mean ICCs across the five datasets would still be considered “poor” according to the Cicchetti and Sparrow (1981) definition.

## 4 Discussion

The present study aimed to demonstrate how connectome-based fingerprint identification accuracy could be used as a simple, intuitive, and sensitive index to adjudicate between functional connectivity-based analysis parameters. Across five small-moderately sized, independent Open Science datasets spanning childhood to adulthood, we found that functional connectome fingerprint identification accuracy ranged from 42% to 77%. We then tested the effect of a range of analysis parameters on ID accuracy and found that GSR, fine-grained (i.e., ∼1,000 ROIs) brain parcellations encompassing cortical regions, number of discriminative edges, and longer scan duration improved identification accuracy — for some datasets to ≥ 90%. Interestingly, pooling fMRI data across datasets to increase sample size decreased, rather than increased, ID accuracy. Importantly, the effects of many of these manipulations on ID accuracy were quantitatively and qualitatively unambiguous, allowing for easy selection of the “best performing” parameters. These “best performing” parameters were then combined into two analysis pipelines that optimize the detection of individual differences in the functional connectome.

Following the investigation of the impact of a set of data acquisition, pre-, and post-processing parameters on fingerprint accuracy, we described two analysis pipelines as follows: (1) Pipeline 1 — GSR, cortical-based parcellation scheme with 1,000 parcels, top 10% of discriminative edges, full scan length, and sample size and; (2) Pipeline 2 — GSR, frontoparietal networks (medial frontal and frontoparietal networks derived from the “Finn” Networks), full scan length, and sample size. With Pipeline 1, accuracies that were 42-77% increased to 68-94%, and with Pipeline 2, they increased to 44-86%. The utility of these pipelines was also supported by the fact that they significantly increased TRT reliability (indexed by ICC) relative to the “standard” pipeline — although it is notable that the ICC values remained poor, according to the traditional categorization (i.e., ICC < 0.4). This echoes the TRT reliability literature which suggests that univariate measures of TRT like ICC for functional connectivity metrics are generally poor (mean: 0.29; 95% CI: 0.23-0.36; Noble et al., 2019). While a direct quantitative comparison of ICC and fingerprint BID is not possible, we can compare them qualitatively given that fingerprint accuracy tracks the ICC values. For example, an ICC = 0.40 implies that the TRT reliability of a functional connectivity measure is fair/moderate, but is above the upper meta-analytic estimate for the ICC across studies (Noble et al., 2019). This means that 40% of the total variance in the measurement of interest is attributed to agreement between the “raters” while the remaining 60% of the variance corresponds to individual variation or measurement errors. In contrast, an ID = 40% would be considered a low level of self-self correlation within a sample. What these comparisons suggest is that, while the ICC values that are typical for the field are considered to reflect “poor” reliability, fingerprint identification accuracy suggests a more optimistic perspective on functional connectome stability. To take one example, BID accuracy for the NYUado dataset processed with Pipeline 1 is 94% (ICC = 0.28), indicating that on average, 23 of 24 participants are correctly identified based on their functional connectomes. Similarly, the BID accuracy derived from Pipeline 1 for the SWU dataset is 90% (ICC = 0.35), which successfully identified, on average, 74 of 82 participants. Hence, connectome-based identification provides a metric of reproducibility of an individual’s functional connectome which is stable on both short-term (days) and long-term (years) from childhood to adulthood. Taken together, these findings mirror the emerging consensus that univariate measures of functional connectivity reliability are generally low whereas multivariate measures (e.g., fingerprinting) tend to be better (Camp et al., 2024; Noble et al., 2017).

Although we found that <u>GSR</u> was beneficial in terms of its impact on fingerprint ID accuracy, it remains the subject of dispute and there is no clear consensus on its application in publicly available pre-processing pipelines (Li et al., 2021a). This disagreement has been traditionally rooted in evidence that showed that GSR introduces negative correlations between brain regions (Saad et al., 2012; Weissenbacher et al., 2009; Murphy et al., 2009), but often countered by its efficacy at mitigating the effects of motion and other sources of noise (Graff et al., 2022; Yan et al., 2013a; Yan et al., 2013b). The debate is ongoing with recent work showing that the global signal could reflect neural information of behavioral relevance: for example, in a sample of 1,094 adults, the topography of the global signal has been shown to capture a positive-negative axis of life outcomes and psychological functions linking to an increased component loading of the frontoparietal network (Li et al., 2019).

Across the lifespan, the topographical patterns of the global signal have revealed spatially specific linear and quadratic associations with age ranging from 6 to 86 years (Nomi et al., 2024). Given the significant and positive effect of GSR on fingerprint accuracy, in combination with strong evidence that GSR reduces the relationships between functional connectivity and head motion, particularly in age groups and vulnerable populations that exhibit heightened in-scanner movement, we recommend its inclusion in processing pipelines.

Describing the functional brain with high-resolution <u>parcellation schemes</u> produced the highest identification accuracies. This likely reflects the fact that increasing the parcel resolution preserves finer properties of the functional connectome from large-scale functional networks to localized brain areas, thus improving the sensitivity and specificity of functional connectivity analyses (Bellec et al., 2015). The choice of selecting a parcel resolution to investigate individual differences in the functional connectome typically depends on the desired spatial resolution of the effects being investigated. We found that <u>cortical regions</u> produced the highest fingerprint accuracy, which is consistent with observations that their functional connectivities are highly reliable compared to that of subcortical and cerebellar structures (Noble et al., 2019; Noble et al., 2017; Shah et al., 2016), which may be driven by factors such as poor signal-to-noise ratio (SNR) due to signal dropout and susceptibility to physiological noise (Noble et al., 2017). It is important to note that there may not be a unique parcellation scheme that captures individual differences in the functional connectome across all brain states (Salehi et al., 2020). Instead, there exists a set of brain parcellations that is condition dependent which provides better parcel definition to detect such individual properties across different states (Salehi et al., 2020). Fingerprints restricted to the <u>medial frontal and frontoparietal networks</u> also yielded the highest fingerprint accuracies compared to the whole-brain connectome. These networks have been shown to exhibit high predictive power and are reproducible in both neurotypical and developing populations (Marek et al., 2019; Jalbrzikowski et al., 2019; Horien et al., 2019; Finn et al., 2015). The frontoparietal network exhibits higher inter-subject variability in its functional connectivity relative to other functional networks including subcortical, limbic, and visual networks (Vanderwal et al. 2017; Mueller et al., 2013), suggesting that this functional network can better discriminate an individual from a group. Restricting the functional connectome fingerprint with the frontoparietal networks (Pipeline 2) would be suitable for some functional connectivity applications but not necessarily for all measures, pipelines, and procedures. Therefore, describing the individual functional connectome with a second fingerprint-informed pipeline that includes a fine-grained parcellation scheme (Schaefer 1000-17Networks; Pipeline 1) offers another optimal strategy to test the reproducibility of functional connectivity applications.

As others have shown (Vanderwal et al., 2021; Vanderwal et al., 2017; Airan et al., 2016; Finn et al., 2015), identification accuracies further increased non-linearly with longer <u>scan duration</u>. Scan duration has long been identified as a factor that affects the reliability of functional connectivity measures (Noble et al., 2019; Elliott et al., 2019; Noble et al., 2017; O’Connor et al., 2017; Mueller et al., 2015; Birn et al., 2013; Van Dijk et al., 2010; Shehzad et al., 2009), along with resting-state conditions such as fixation, eyes-opened, and eyes-closed (Wang et al., 2017; Zuo et al., 2015; Tagliazucchi and Laufs, 2014; Patriat et al., 2013). While initial reports suggested that acquisition times of ∼5 min are sufficient to yield fair-to-excellent reliability (Van Dijk et al., 2010; Shehzad et al., 2009), more recent studies have suggested that much longer duration scans are required to yield the most reliable (e.g., > 36 min; Noble et al., 2017) and reproducible (e.g., 15-25 min; Anderson et al., 2011) metrics. Researchers have employed several strategies to increase the amount of fMRI data available per participant, including longer scan duration (Noble et al., 2019; Noble et al., 2017; Tomasi et al., 2016; Shah et al., 2016), repeated scans within-/between-sessions (Cho et al., 2020; Elliott et al., 2019; O’Connor et al., 2017; Noble et al., 2017), and higher sampling rates (Birn et al., 2013). Acquiring and combining more scans for each individual across different brain states could further improve the fingerprint accuracy as within-subject variability may be better captured than at rest (Finn et al., 2017). This echoes recent evidence which suggests that having more fMRI data per participant increases the ability to capture unique properties of the functional connectome using resting-state and naturalistic conditions (Vanderwal et al., 2017). It is, however, challenging to identify an ideal amount of fMRI data that is needed per participant to optimize the reliability of functional connectivity — longer scan lengths undermine the practical utility of fMRI, especially in developing and clinical populations with lower in-scanner tolerance or wakefulness.

In agreement with previous work (Vanderwal et al., 2021; Vanderwal et al., 2017), identification patterns were improved when a greater number of <u>discriminative edges</u> was used to compute a given individual’s functional connectivity profiles. This finding showed that as few as 10% of the most discriminative edges were needed to yield good fingerprint accuracies as there was no significant difference in identification accuracies for the top 10% or 75% of the most discriminative edges relative to the full whole-brain connectome. This suggests that computational complexity and resource demands could be used by downsizing the functional connectome to the top 10% of discriminative edges. This also suggests that there are some edges that do not contribute to the identifiability of an individual’s functional connectome, thus positing that the entire functional connectome may not be required to compute functional connectome fingerprints.

We also pooled fMRI data across the five datasets to investigate the impact of <u>sample size</u>, but this did not improve the fingerprint accuracies. That is, larger sample sizes (within the ranges tested here) seem to increase the difficulty of differentiating an individual from a group. Similar observations have been reported which showed that fingerprint accuracy decreases non-linearly when the sample size derived from the Human Connectome Project increases from typical (e.g., *N* = 100) to large (e.g., *N* = 1,000) (Li et al., 2021; Waller et al., 2017). A potential limitation of this sampling strategy is that we aggregated the fMRI data across a wide range of age groups (i.e., 7-43 years) which may not be ideal to detect individual differences in the functional connectome. There could also be a site effect that requires further investigation, particularly in a larger dataset acquired from several more sites. This may be a challenge as a downside of the connectome-based identification procedure is that a minimum of two imaging scans is required which is not always feasible during cross-sectional study designs. Future work may investigate the utility of split-half fingerprinting as a proxy method (King et al., 2023; Wang et al., 2021). Along the same lines, employing a larger multisite dataset could allow the investigation of inter-scan intervals on fingerprint accuracy. The CoRR datasets used in this study make the comparison very difficult due to the wide range of inter-scan intervals from days to years.

Although we aimed to identify a pipeline that would optimize functional connectome fingerprint ID, and while high identification accuracies were attained for some samples, the accuracy did not reach 100% for any sample. This demonstrates that there are some individuals who remain incorrectly identified even with the fingerprint-informed pipelines. This suggests that there is scope to identify further factors that might influence fingerprint identification accuracy. On the other hand, it may also suggest that there are individuals who remain misidentified (Greene et al., 2022), even in the context of overall high ID accuracy. We attempted to quantify the degree of misidentification for a given individual from a group using the RR metric which was as low as 2% when the optimal pipelines were employed. The RR metric did not provide additional discriminative information beyond BID accuracy, however.

Can we expect resting-state functional connectome fingerprint ID accuracy to be perfect? There is evidence for 100% identification accuracy, achieved using naturalistic paradigms, albeit in a small sample (Vanderwal et al., 2017). The answer may lie in the cognitive states of participants being scanned (Finn and Rosenberg, 2021; Finn et al., 2017); scanning conditions that evoke more specific cognitive states tend to be associated with higher fingerprint accuracy relative to rest (Vanderwal et al., 2017; Finn et al., 2017). It is possible that connectome-based identification accuracy at rest may reflect the influence of arousal, thought, and even physiological processes (Finn et al., 2017). Another important consideration is the influence of anatomical variability on individual differences in functional connectivity estimates that have been characterized as a mixture of morphological and functional variability (Langs et al., 2015). Converging evidence from human and primate studies have shown that there is a significant relationship between structural and functional connectivities such that structural connectivity may predict inter-individual variability in cognition (Sarwar et al., 2020; Sporns, 2013). This relationship may, however, be heterogeneous as the structure-function coupling could be stronger for some networks but weaker for others (Gu et al., 2021; Suárez et al., 2020; Baum et al., 2019). Future work could investigate the contributions of anatomical variability (e.g., morphology) in measuring individual differences in functional connectome identifiability. Yet another avenue could focus on investigating whether unique or similar structure-function coupling across individuals relates to better or worse connectome-based identification.

## 5 Conclusion

In this study, we showed that functional connectome fingerprint accuracy can be used as a simple, intuitive, and sensitive metric for examining the effects of different functional connectivity analysis parameters in smaller and moderately sized samples, and in both developmental and adult datasets. By examining a set of data acquisition, pre-, and post-processing parameters, we identified a number of recommended steps that form two pipelines optimized for reproducible and stable functional connectivity at the individual level (**Table 3**). The pipelines comprise the following strategies: (I) applying GSR as a denoising strategy; (II) fine-grained brain parcellations; (III) cortical regions relative to subcortical and cerebellar structures; (IV) predictive functional networks such as medial frontal and frontoparietal networks; (V) discriminative edges; and (VI) increasing scan duration. In contrast, increasing the sample size by pooling resting-state fMRI data across different populations do not necessarily yield higher fingerprint accuracies. We suggest that the inclusion of fingerprint accuracy in analysis workflows can help researchers identify the most reproducible pipelines and as a result, minimize the “researcher degrees of freedom” to ultimately achieve robust and reproducible brain-behavior relationships for biomarker discovery.

**Table 3.** Recommendations for describing an optimal pipeline to reliably measure the functional connectome at the individual level. The recommendations follow the underlying factors that influence fingerprint accuracy at the data pre- and post-processing stages as well as during the data acquisition phase.

| Recommendations |  |
| --- | --- |
| Factors | Influence on fingerprint accuracy |
| **GSR** | **Increases** by regressing out the GS in the fMRI timeseries |
| **Parcellation Scheme** | **Increases** with higher resolutions (Shen 268 → Schaefer 1000) |
| **Parcel Resolution** | **Increases** as parcel resolution increases (7 → 1095) |
| **Parcel Coverage** | **Increases** with cortical regions<br>**Decreases** with subcortical and cerebellar structures |
| **Network Organisation** | **Increases** with medial frontal and frontoparietal networks<br>**Increases** with medial frontal + frontoparietal networks<br>**Decreases** with visual, subcortical and limbic networks |
| **Discriminative Edges** | **Increases** with higher number of differentiating edges (top 10%)<br>**Decreases** with lower number of differentiating edges (top 1%) |
| **Scan Duration** | **Increases** with longer scan lengths (0.5 min → full scan duration) |
| **Amount of fMRI Data** | **Decreases** with increasing sample sizes (20 → 264) |

## ACKNOWLEDGMENTS

We kindly thank the Consortium for Replicability and Reproducibility for allowing the neuroimaging data to be publicly accessible.

## Authorship Contribution Statement

**Jivesh Ramduny:** Conceptualization, Methodology, Formal analysis, Investigation, Writing — original draft, review & editing. **Tamara Vanderwal:** Conceptualization, Investigation, Writing — original draft, review & editing. **Clare Kelly:** Conceptualization, Funding, Methodology, Formal analysis, Investigation, Writing — original draft, review & editing, Supervision, Project administration.

## References

Airan, R. D., Vogelstein, J. T., Pillai, J. J., Caffo, B., Pekar, J. J., & Sair, H. I. (2016). Factors affecting characterization and localization of interindividual differences in functional connectivity using MRI. Human Brain Mapping, 37(5), 1986–1997.

Anderson, J. S., Ferguson, M. A., Lopez-Larson, M., & Yurgelun-Todd, D. (2011). Reproducibility of single-subject functional connectivity measurements. American Journal of Neuroradiology, 32(3), 548–555.

Bartko, J. J. (1976). On various intraclass correlation reliability coefficients. Psychological Bulletin, 83(5), 762–765.

Baum, G. L., Cui, Z., Roalf, D. R., Ciric, R., Betzel, R. F., Larsen, B. et al. (2019). Development of structure–function coupling in human brain networks during Youth. Proceedings of the National Academy of Sciences, 117(1), 771–778.

Bellec, P., Benhajali, Y., Carbonell, F., Dansereau, C., Albouy, G., Pelland, M. et al. (2015). Impact of the resolution of brain parcels on Connectome-wide association studies in fmri. NeuroImage, 123, 212–228.

Birn, R. M., Molloy, E. K., Patriat, R., Parker, T., Meier, T. B., Kirk, G. R. et al. (2013). The effect of scan length on the reliability of resting-state fmri connectivity estimates. NeuroImage, 83, 550–558.

Byrge, L., & Kennedy, D. P. (2019). High-accuracy individual identification using a “thin slice” of the functional connectome. Network Neuroscience, 3(2), 363–383.

Cai, B., Zhang, G., Zhang, A., Xiao, L., Hu, W., Stephen, J. M. et al. (2021). Functional connectome fingerprinting: Identifying individuals and predicting cognitive functions via autoencoder. Human Brain Mapping, 42(9), 2691–2705.

Camp, C. C., Noble, S., Scheinost, D., Stringaris, A., & Nielson, D. M. (2024). Test-retest reliability of functional connectivity in adolescents with depression. Biological Psychiatry: Cognitive Neuroscience and Neuroimaging, 9(1), 21–29.

Castellanos, F. X., Di Martino, A., Craddock, R. C., Mehta, A. D., & Milham, M. P. (2013). Clinical applications of the functional connectome. NeuroImage, 80, 527–540.

Chen, G., Taylor, P. A., Haller, S. P., Kircanski, K., Stoddard, J., Pine, D. S. et al. (2018). Intraclass correlation: Improved modeling approaches and applications for neuroimaging. Human Brain Mapping, 39(3), 1187–1206.

Cho, J. W., Korchmaros, A., Vogelstein, J. T., Milham, M., & Xu, T. (2020). Impact of concatenating fmri data on reliability for Functional Connectomics. NeuroImage, 226, 117549.

Cicchetti, D. V., & Sparrow S. A. (1981). Developing criteria for establishing interrater reliability of specific items: applications to assessment of adaptive behavior. American journal of mental deficiency, 86(2), 127–137.

Ciric, R., Wolf, D. H., Power, J. D., Roalf, D. R., Baum, G. L., Ruparel, K. et al. (2017). Benchmarking of participant-level confound regression strategies for the control of motion artifact in studies of Functional Connectivity. NeuroImage, 154, 174–187.

Craddock, C., Benhajali, Y., Chu, C., Chouinard, F., Evans, A., Jakab, A. et al. (2013). The Neuro Bureau Preprocessing Initiative: open sharing of preprocessed neuroimaging data and derivatives. Front. Neuroinform. Conference Abstract: Neuroinformatics.

Dufford, A. J., Noble, S., Gao, S., & Scheinost, D. (2021). The instability of functional connectomes across the first year of life. Developmental Cognitive Neuroscience, 51, 101007.

Elliott, M. L., Knodt, A. R., Cooke, M., Kim, M. J., Melzer, T. R., Keenan, R. et al. (2019). General functional connectivity: Shared features of resting-state and task fmri drive reliable and heritable individual differences in Functional Brain Networks. NeuroImage, 189, 516–532.

Finn, E. S., & Rosenberg, M. D. (2021). Beyond fingerprinting: Choosing predictive connectomes over reliable connectomes. NeuroImage, 239, 118254.

Finn, E. S., Scheinost, D., Finn, D. M., Shen, X., Papademetris, X., & Constable, R. T. (2017). Can brain state be manipulated to emphasize individual differences in functional connectivity? NeuroImage, 160, 140–151.

Finn, E. S., Shen, X., Scheinost, D., Rosenberg, M. D., Huang, J., Chun, M. M. et al. (2015). Functional connectome fingerprinting: Identifying individuals using patterns of brain connectivity. Nature Neuroscience, 18(11), 1664–1671.

Fox, M. D., Zhang, D., Snyder, A. Z., & Raichle, M. E. (2009). The global signal and observed anticorrelated resting State Brain Networks. Journal of Neurophysiology, 101(6), 3270–3283.

Glasser, M. F., Coalson, T. S., Robinson, E. C., Hacker, C. D., Harwell, J., Yacoub, E. et al. (2016). A multi-modal parcellation of human cerebral cortex. Nature, 536(7615), 171–178.

Graff, K., Tansey, R., Ip, A., Rohr, C., Dimond, D., Dewey, D., & Bray, S. (2022). Benchmarking common preprocessing strategies in early childhood functional connectivity and intersubject correlation fmri. Developmental Cognitive Neuroscience, 54, 101087.

Gratton, C., Kraus, B. T., Greene, D. J., Gordon, E. M., Laumann, T. O., Nelson, S. M. et al. (2020). Defining individual-specific functional neuroanatomy for precision psychiatry. Biological Psychiatry, 88(1), 28–39.

Greene, A. S., Shen, X., Noble, S., Horien, C., Hahn, C. A., Arora, J. et al. (2022). Brain–phenotype models fail for individuals who defy sample stereotypes. Nature, 609(7925), 109–118.

Greve, D. N., & Fischl, B. (2009). Accurate and robust brain image alignment using boundary-based registration. NeuroImage, 48(1), 63–72.

Gu, Z., Jamison, K. W., Sabuncu, M. R., & Kuceyeski, A. (2021). Heritability and interindividual variability of regional structure-function coupling. Nature Communications, 12(1).

Heale, R., & Twycross, A. (2015). Validity and reliability in quantitative studies. Evidence Based Nursing, 18(3), 66–67.

Horien, C., Shen, X., Scheinost, D., & Constable, R. T. (2019). The individual functional connectome is unique and stable over months to years. NeuroImage, 189, 676–687.

Jalbrzikowski, M., Liu, F., Foran, W., Klei, L., Calabro, F. J., Roeder, K. et al. (2019). Functional connectome fingerprinting accuracy in youths and adults is similar when examined on the same day and 1.5-years apart. Human Brain Mapping, 41(15), 4187–4199.

Jenkinson, M., Bannister, P., Brady, M., & Smith, S. (2002). Improved optimization for the robust and accurate linear registration and motion correction of brain images. NeuroImage, 17(2), 825–841.

Kaufmann, T., Alnæs, D., Doan, N. T., Brandt, C. L., Andreassen, O. A., & Westlye, L. T. (2017). Delayed stabilization and individualization in connectome development are related to psychiatric disorders. Nature Neuroscience, 20(4), 513–515.

King, G., Truzzi, A., & Cusack, R. (2023). The confound of head position in within-session connectome fingerprinting in infants. NeuroImage, 265, 119808.

Koo, T. K., & Li, M. Y. (2016). A guideline of selecting and reporting intraclass correlation coefficients for Reliability Research. Journal of Chiropractic Medicine, 15(2), 155–163.

Langs, G., Wang, D., Golland, P., Mueller, S., Pan, R., Sabuncu, M. R. et al. (2015). Identifying shared brain networks in individuals by decoupling functional and anatomical variability. Cerebral Cortex, 26(10), 4004–4014.

Li, J., Bolt, T., Bzdok, D., Nomi, J. S., Yeo, B. T., Spreng, R. N., & Uddin, L. Q. (2019). Topography and behavioral relevance of the global signal in the human brain. Scientific Reports, 9(1).

Li, X., Esper, N. B., Ai, L., Giavasis, S., Jin, H., Feczko, E. et al. (2021a). Moving beyond processing and analysis-related variation in Neuroscience. bioRxiv.

Li, K., Wisner, K., & Atluri, G. (2021b). Feature selection framework for functional connectome fingerprinting. Human Brain Mapping, 42(12), 3717–3732.

Liljequist, D., Elfving, B., & Skavberg Roaldsen, K. (2019). Intraclass correlation – a discussion and demonstration of basic features. PLOS ONE, 14(7).

Litwińczuk, M. C., Muhlert, N., Cloutman, L., Trujillo-Barreto, N., & Woollams, A. (2022). Combination of structural and functional connectivity explains unique variation in specific domains of cognitive function. NeuroImage, 262, 119531.

Marek, S., Tervo-Clemmens, B., Nielsen, A. N., Wheelock, M. D., Miller, R. L., Laumann, T. O. et al. (2019). Identifying reproducible individual differences in childhood functional brain networks: An ABCD study. Developmental Cognitive Neuroscience, 40, 100706.

Margulies, D. S., Vincent, J. L., Kelly, C., Lohmann, G., Uddin, L. Q., Biswal, B. B. et al. (2009). Precuneus shares intrinsic functional architecture in humans and monkeys. Proceedings of the National Academy of Sciences, 106(47), 20069–20074.

Mellinger, C. D., & Hanson, T. A. (2021). Methodological Considerations for survey research: Validity, reliability, and Quantitative Analysis. Linguistica Antverpiensia, New Series – Themes in Translation Studies, 19, 172–190.

Mueller, S., Wang, D., Fox, M. D., Pan, R., Lu, J., Li, K. et al. (2015). Reliability Correction for functional connectivity: Theory and implementation. Human Brain Mapping, 36(11), 4664–4680.

Mueller, S., Wang, D., Fox, M. D., Yeo, B. T. T., Sepulcre, J., Sabuncu, M. R. et al. (2013). Individual variability in functional connectivity architecture of the human brain. Neuron, 77(3), 586–595.

Murphy, K., Birn, R. M., Handwerker, D. A., Jones, T. B., & Bandettini, P. A. (2009). The impact of global signal regression on resting state correlations: Are anti-correlated networks introduced? NeuroImage, 44(3), 893–905.

Noble, S., Scheinost, D., & Constable, R. T. (2019). A decade of test-retest reliability of functional connectivity: A systematic review and meta-analysis. NeuroImage, 203, 116157.

Noble, S., Scheinost, D., & Constable, R. T. (2021). A guide to the measurement and interpretation of fmri test-retest reliability. Current Opinion in Behavioral Sciences, 40, 27–32.

Noble, S., Spann, M. N., Tokoglu, F., Shen, X., Constable, R. T., & Scheinost, D. (2017). Influences on the test–retest reliability of functional connectivity MRI and its relationship with behavioral utility. Cerebral Cortex, 27(11), 5415–5429.

Nomi, J. S., Bzdok, D., Li, J., Bolt, T., Chang, C., Kornfeld, S. et al. (2024). Systematic cross-sectional age-associations in global fmri signal topography. Imaging Neuroscience, 2, 1–13.

O’Connor, D., Potler, N. V., Kovacs, M., Xu, T., Ai, L., Pellman, J. et al. (2017). The Healthy Brain Network Serial scanning initiative: A resource for evaluating inter-individual differences and their reliabilities across scan conditions and sessions. GigaScience, 6(2).

Parkes, L., Fulcher, B., Yücel, M., & Fornito, A. (2018). An evaluation of the efficacy, reliability, and sensitivity of motion correction strategies for resting-state functional MRI. NeuroImage, 171, 415–436.

Patriat, R., Molloy, E. K., Meier, T. B., Kirk, G. R., Nair, V. A., Meyerand, M. E. et al. (2013). The effect of resting condition on resting-state fmri reliability and consistency: A comparison between resting with eyes open, closed, and fixated. NeuroImage, 78, 463–473.

Power, J. D., Mitra, A., Laumann, T. O., Snyder, A. Z., Schlaggar, B. L., & Petersen, S. E. (2014). Methods to detect, characterize, and remove motion artifact in resting state fmri. NeuroImage, 84, 320–341.

Power, J. D., Plitt, M., Kundu, P., Bandettini, P. A., & Martin, A. (2017). Temporal interpolation alters motion in fmri scans: Magnitudes and consequences for artifact detection. PLOS ONE, 12(9).

Saad, Z. S., Gotts, S. J., Murphy, K., Chen, G., Jo, H. J., Martin, A., & Cox, R. W. (2012). Trouble at rest: How correlation patterns and group differences become distorted after global signal regression. Brain Connectivity, 2(1), 25–32.

Salehi, M., Greene, A. S., Karbasi, A., Shen, X., Scheinost, D., & Constable, R. T. (2020). There is no single functional atlas even for a single individual: Functional parcel definitions change with Task. NeuroImage, 208, 116366.

Sarwar, T., Tian, Y., Yeo, B. T. T., Ramamohanarao, K., & Zalesky, A. (2020). Structure-function coupling in the human connectome: A machine learning approach. NeuroImage, 226, 117609.

Satterthwaite, T. D., Ciric, R., Roalf, D. R., Davatzikos, C., Bassett, D. S., & Wolf, D. H. (2017). Motion artifact in studies of functional connectivity: Characteristics and mitigation strategies. Human Brain Mapping, 40(7), 2033–2051.

Schaefer, A., Kong, R., Gordon, E. M., Laumann, T. O., Zuo, X.-N., Holmes, A. J. et al. (2018). Local-global parcellation of the human cerebral cortex from intrinsic functional connectivity MRI. Cerebral Cortex, 28(9), 3095–3114.

Shah, L. M., Cramer, J. A., Ferguson, M. A., Birn, R. M., & Anderson, J. S. (2016). Reliability and reproducibility of individual differences in functional connectivity acquired during task and resting state. Brain and Behavior, 6(5).

Shehzad, Z., Kelly, A. M., Reiss, P. T., Gee, D. G., Gotimer, K., Uddin, L. Q. et al. (2009). The Resting Brain: Unconstrained yet reliable. Cerebral Cortex, 19(10), 2209–2229.

Shen, X., Tokoglu, F., Papademetris, X., & Constable, R. T. (2013). Groupwise whole-brain parcellation from resting-state fmri data for network node identification. NeuroImage, 82, 403–415.

Shrout, P. E., & Fleiss, J. L. (1979). Intraclass correlations: Uses in assessing rater reliability. Psychological Bulletin, 86(2), 420–428.

Sporns, O. (2013). Structure and function of Complex Brain Networks. Dialogues in Clinical Neuroscience, 15(3), 247–262.

St-Onge, F., Javanray, M., Pichet Binette, A., Strikwerda-Brown, C., Remz, J., Spreng, R. N. et al. (2023). Functional connectome fingerprinting across the lifespan. Network Neuroscience, 7, 1206–27.

Suárez, L. E., Markello, R. D., Betzel, R. F., & Misic, B. (2020). Linking structure and function in Macroscale Brain Networks. Trends in Cognitive Sciences, 24(4), 302–315.

Tagliazucchi, E., & Laufs, H. (2014). Decoding wakefulness levels from typical fmri resting-state data reveals reliable drifts between wakefulness and sleep. Neuron, 82(3), 695–708.

Tomasi, D. G., Shokri-Kojori, E., & Volkow, N. D. (2016). Temporal evolution of brain functional connectivity metrics: Could 7 min of rest be enough? Cerebral Cortex, 27, 4153–4165.

Tozzi, L., Fleming, S. L., Taylor, Z. D., Raterink, C. D., & Williams, L. M. (2020). Test-retest reliability of the human functional connectome over consecutive days: Identifying highly reliable portions and assessing the impact of methodological choices. Network Neuroscience, 4(3), 925–945.

Urchs, S., Armoza, J., Benhajali, Y., St-Aubin, J., Orban, P., & Bellec, P. (2017). Mist: A multi-resolution parcellation of Functional Brain Networks. MNI Open Research, 1, 3.

Van Dijk, K. R., Hedden, T., Venkataraman, A., Evans, K. C., Lazar, S. W., & Buckner, R. L. (2010). Intrinsic functional connectivity as a tool for human Connectomics: Theory, properties, and Optimization. Journal of Neurophysiology, 103(1), 297–321.

Vanderwal, T., Eilbott, J., Finn, E. S., Craddock, R. C., Turnbull, A., & Castellanos, F. X. (2017). Individual differences in functional connectivity during naturalistic viewing conditions. NeuroImage, 157, 521–530.

Vanderwal, T., Eilbott, J., Kelly, C., Frew, S. R., Woodward, T. S., Milham, M. P., & Castellanos, F. X. (2021). Stability and similarity of the pediatric connectome as developmental measures. NeuroImage, 226, 117537.

Waller, L., Walter, H., Kruschwitz, J. D., Reuter, L., Müller, S., Erk, S., & Veer, I. M. (2017). Evaluating the replicability, specificity, and generalizability of connectome fingerprints. NeuroImage, 158, 371–377.

Wang, J., Han, J., Nguyen, V. T., Guo, L., & Guo, C. C. (2017). Improving the test-retest reliability of resting state fmri by removing the impact of sleep. Frontiers in Neuroscience, 11.

Wang, Q., Xu, Y., Zhao, T., Xu, Z., He, Y., & Liao, X. (2021). Individual uniqueness in the neonatal functional connectome. Cerebral Cortex, 31(8), 3701–3712.

Weissenbacher, A., Kasess, C., Gerstl, F., Lanzenberger, R., Moser, E., & Windischberger, C. (2009). Correlations and anticorrelations in resting-state functional connectivity MRI: A quantitative comparison of preprocessing strategies. NeuroImage, 47(4), 1408–1416.

Yan, C.-G., Cheung, B., Kelly, C., Colcombe, S., Craddock, R. C., Di Martino, A. et al. (2013a). A comprehensive assessment of regional variation in the impact of head micromovements on Functional connectomics. NeuroImage, 76, 183–201.

Yan, C.-G., Craddock, R. C., Zuo, X.-N., Zang,Y.-F., & Milham, M. P. (2013b). Standardizing the intrinsic brain: Towards robust measurement of inter-individual variation in 1000 functional connectomes. NeuroImage, 80, 246–262.

Yeo, B. T. T., Krienen, F. M., Sepulcre, J., Sabuncu, M. R., Lashkari, D., Hollinshead, M. et al. (2011). The organization of the human cerebral cortex estimated by intrinsic functional connectivity. Journal of Neurophysiology, 106(3), 1125–1165.

Zuo, X.-N, Anderson, J. S., Bellec, P., Birn, R. M., Biswal, B. B., Blautzik, J. et al. (2014). An open science resource for establishing reliability and reproducibility in functional connectomics. Sci Data 1, 140049.

Zuo, X.-N., Biswal, B. B., & Poldrack, R. A. (2019). Editorial: Reliability and reproducibility in functional connectomics. Frontiers in Neuroscience, 13.

Zou, Q., Miao, X., Liu, D., Wang, D. J. J., Zhuo, Y., & Gao, J.-H. (2015). Reliability comparison of spontaneous brain activities between Bold and CBF contrasts in eyes-open and eyes-closed resting states. NeuroImage, 121, 91–105.

